# Live Monitoring of Inflammation Reveals Tissue and Sex-specific Responses to Western Diet and Butyrate treatment

**DOI:** 10.1101/2021.09.22.461384

**Authors:** Raiza Bonomo, Sarah Talley, Jomana Hatahet, Chaitanya Gavini, Tyler Cook, Ben Chun, Pete Kekenes-Huskey, Gregory Aubert, Edward Campbell, Virginie Mansuy-Aubert

## Abstract

Obesity is a current epidemic, affecting millions of individuals worldwide. Chronic obesity is characterized by a low-grade systemic inflammation besides not being a classic inflammatory disease. Many studies have tried to identify inflammatory insults dysregulated by a Westernized diet – consisted of high fat, high sucrose, and high cholesterol –mainly focusing on production and secretion of inflammatory cytokines. The gut microbiome and derived metabolites, including the short-chain fatty acid butyrate, have received increased attention as underlying some of the obesogenic features. In the present work, we utilized a novel biosensor mouse model capable of monitoring in vivo inflammation. We observed tissue- and sex- specific caspase-1 activation patterns in obese mice and treated with butyrate. Our work utilizing a caspase-1 biosensor mouse model, flow cytometry and computational analyses and offers new mechanistic insights underlying the effect of butyrate in obesity and its complications.

## Introduction

Obesity (body-mass index ≥30kg/m^2^) is the result of long-term energy homeostasis imbalance (e.g increased food intake, decreased energy expenditure) involving brain and peripheral organs (1). Obesity is the cause of many complications, including diabetes, liver diseases, cardiovascular diseases or even neurological dysfunction such as memory dysfunction or pain (2, 3)

Recently, the gut microbiome and its derivative metabolites have received attention as potential causes and/or consequences of obesity (4–14). A seminal work by Backhed et al. showed that germ-free mice are protected from developing diet-induced obesity, demonstrating the crucial role played by gut microbiome in energy balance, substrate utilization and glucose homeostasis (15). A novel study by Mocanu et al. revealed that a diet rich in fermentable fibers in conjunction with a single-dose fecal transplantation from lean donors to obese patients improved insulin sensitivity (16). This work demonstrated that modulation of gut microbiome holds therapeutic promise in alleviating obesity-associated comorbidities. The microbiome produces byproducts from fiber fermentation, including short-chain fatty acids: butyrate, propionate, acetate. We and others have showed that WD led to drastic alterations of gut microbiome composition, decreasing the relative abundance of butyrate-producers (17). Alterations of gut microbiome and specifically the decrease in butyrate-producing species are commonly observed in obesity (6). Butyrate supplementation improves some complications of obesity including glucose intolerance (6, 11, 12, 18), heart disease (19), and pain (17, 20). However, no molecular mechanisms underlying butyrate’s effects have clearly been identified in obese models. Reports have demonstrated that butyrate has anti-inflammatory effects regulating T cell function, macrophage polarization and act as antimicrobial (11, 20–25). Particularly, butyrate decreases the production of inflammatory cytokines, including IL-1β, in adipose tissue of genetic model of obesity (*db/db*) (26). Low-grade inflammation is thought to play an important role in the onset and progression of obesity and its complications (27–29) but the link between butyrate and inflammation in obesity is unclear. In the present work, we took advantage of a recent engineered mouse model to longitudinally monitor live caspase-1 mediated inflammation *in vivo* in male and female obese mice. We quantified caspase-1 biosensor activation upon WD feeding and treatment with butyrate in tissues involved in obesity and its complications including brain, heart, pancreas, white and brown adipose tissue, liver, intestine. We observed a sex- and tissue-specific inflammasome activation upon WD nutrition and butyrate treatment.

## Results

### Caspase-1 biosensor mice display expected WD-induced phenotype

We used the caspase-1 biosensor model that has been recently generated to monitor inflammation in vivo in a spatiotemporal manner (30). Briefly, transgenic mice express a circularly permuted form of luciferase that becomes bioluminescent in response to active caspase-1, allowing the visualization of caspase-1-mediated inflammatory response in live animals (30). To validate the use of this model, we first confirmed that the genetic manipulation did not affect whole body metabolism and response to WD. Biosensor mice were placed on NC or WD feeding paradigms for a total of 20 weeks. We measured metabolic parameters longitudinally. As expected, we noted expected WD-induced metabolic syndrome features such as increased body weight in comparison to NC-fed mice (Figure 1, A), accompanied by WD induced glucose intolerance (Figure 1, B–C). WD-fed biosensor animals also presented elevated fasting glucose levels (Figure 1, D) after 17 weeks on the feeding paradigm. We also confirmed an increase in energy expenditure (Figure 1, E) and decreased respiratory exchange ratio (RER (Figure 1, F), with no changes in cumulative food consumption (Figure 1, G) when the mice were switched from a NC to WD. Our data demonstrate that the caspase-1 biosensor mouse model show expected behavioral features upon prolonged WD feeding in males and females.

**Figure 1.**
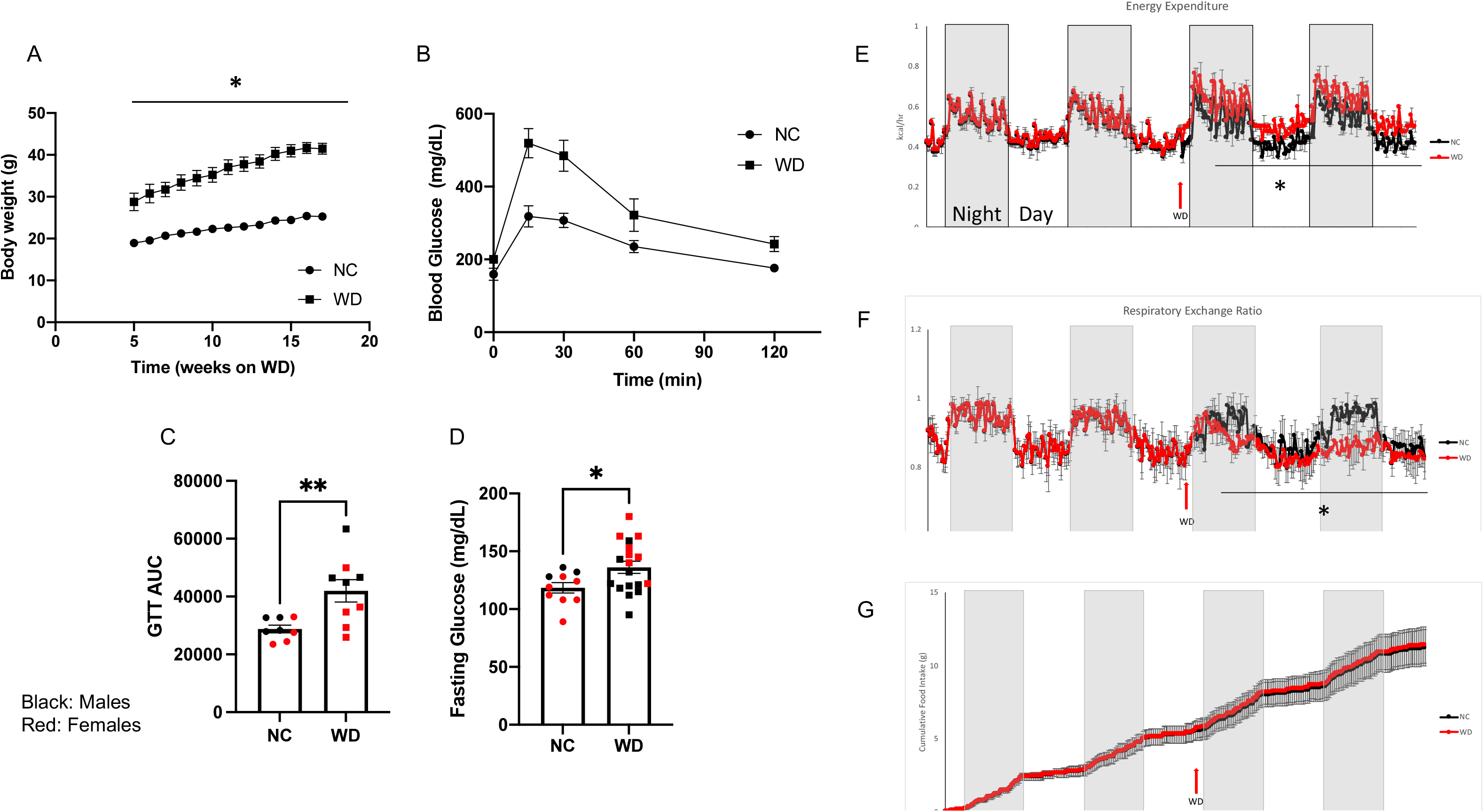
Caspase-1 Biosensor Mice Display Expected WD-induced Phenotype. (A) Body weight of normal chow (NC) and Western diet (WD)-fed mice. (B) Glucose tolerance test (GTT) between NC- and WD-fed mice. (C) Representative GTT area under the curve (AUC). (D) Overnight fasting glucose blood levels. (E) Energy expenditure, (F) Respiratory exchange ratio, and (G) Cumulative food intake from NC and WD-fed biosensor mice. Student’s t test between NC and WD. All values represent mean±S.E.M, *p<0.05, n=4-8 mice/group; males and females.

### Obese mice exhibit gradual increase in whole body caspase-1 activation in vivo

Concurrent with phenotypic analyses, we performed longitudinal imaging of NC- and WD-fed biosensor mice. Males and females were imaged using in vivo imaging system (IVIS). Obese animals from both sexes showed increased bioluminescence signal when compared to lean mice as seen in the representative images in Figure 2, A. Both, male and female WD-fed mice, showed a rise in signal from week 6 to week 10 that was sustained at week 17 of WD feeding paradigm, although obese male mice exhibited stronger bioluminescence relative to female mice (Figure 2 B–C). Together, our results indicate that WD lead to activation of the engineered caspase-1 biosensor in males and females over time with a slight sexual dimorphism.

**Figure 2.**
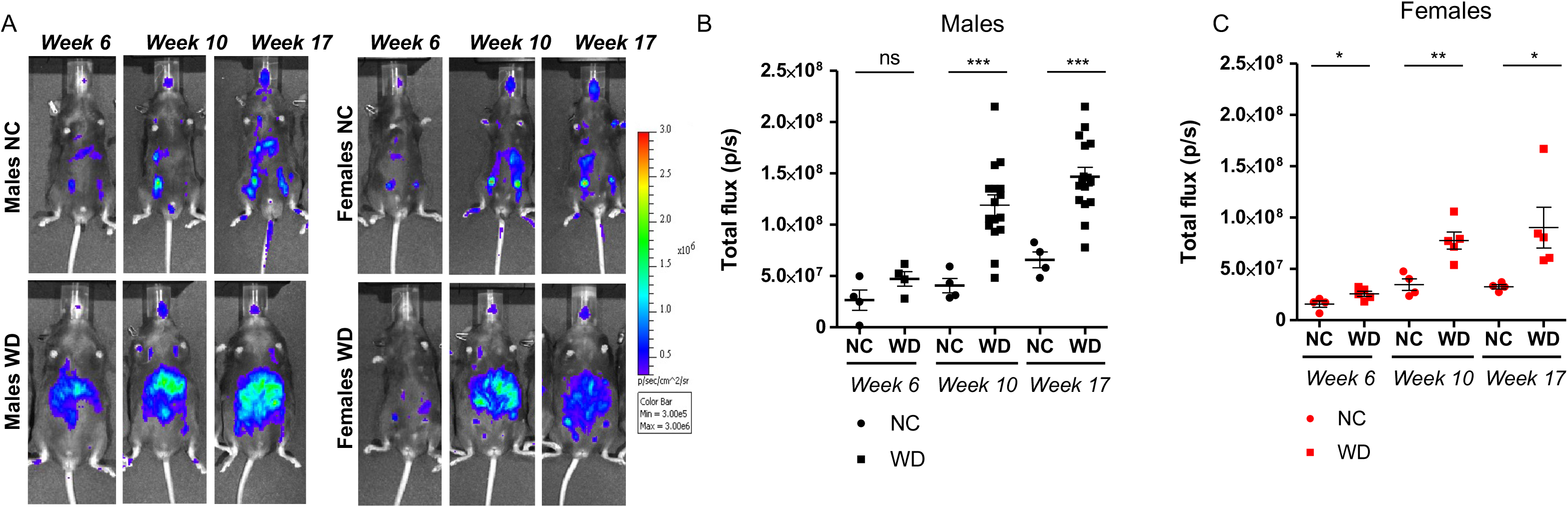
Obese Mice Exhibit Gradual Increase in Whole Body Caspase-1 Activation In Vivo. Representative IVIS images of normal chow (NC) and Western diet (WD)-fed males and females over time. (B) and (C) Bioluminescence signal quantification for males and females. Student’s t test between NC and WD. All values represent mean±S.E.M, *p<0.05; n=4-8 mice/group; males and females.

### Tributyrin and WD alter caspase-1 activation differentially in tissues from male and female mice

Numerous studies have demonstrated that long-term butyrate treatment is protective in mouse models of diet-induced obesity, reducing body weight gain and insulin resistance in obese mice (31, 32) and attenuating obesity-associated inflammation. To better understand the early mechanistic changes driving these protective effects, we aimed to develop a paradigm of short-term butyrate administration that would allow us to assess the inflammatory changes occurring in obese mice prior to butyrate-induced weight loss and improved fasting glucose. We tested whether an increase in circulating butyrate would modify caspase-1 activation and immune cells in vivo in CNS and metabolic tissues. First, we defined the dose and time-course of tributyrin administration. In a dose response pilot study, using serum lipidomics, we observed that 5g/kg of tributyrin led to 25 times increase in circulating butyrate after 2 hours, persisting until 48h post-treatments (not shown). We established a treatment paradigm by administrating 5g/kg of tributyrin every 48h to the biosensor mouse model for two weeks (Figure 3, A). Two weeks of tributyrin trended towards attenuating fasting glucose levels of WD-fed mice (Figure 3, B) without an effect on body weight as previously observed (Figure 3, C). Two weeks of tributyrin administration also did not significantly reduce systemic biosensor activation *in vivo* (supplemental Fig. 1).

**Figure 3.**
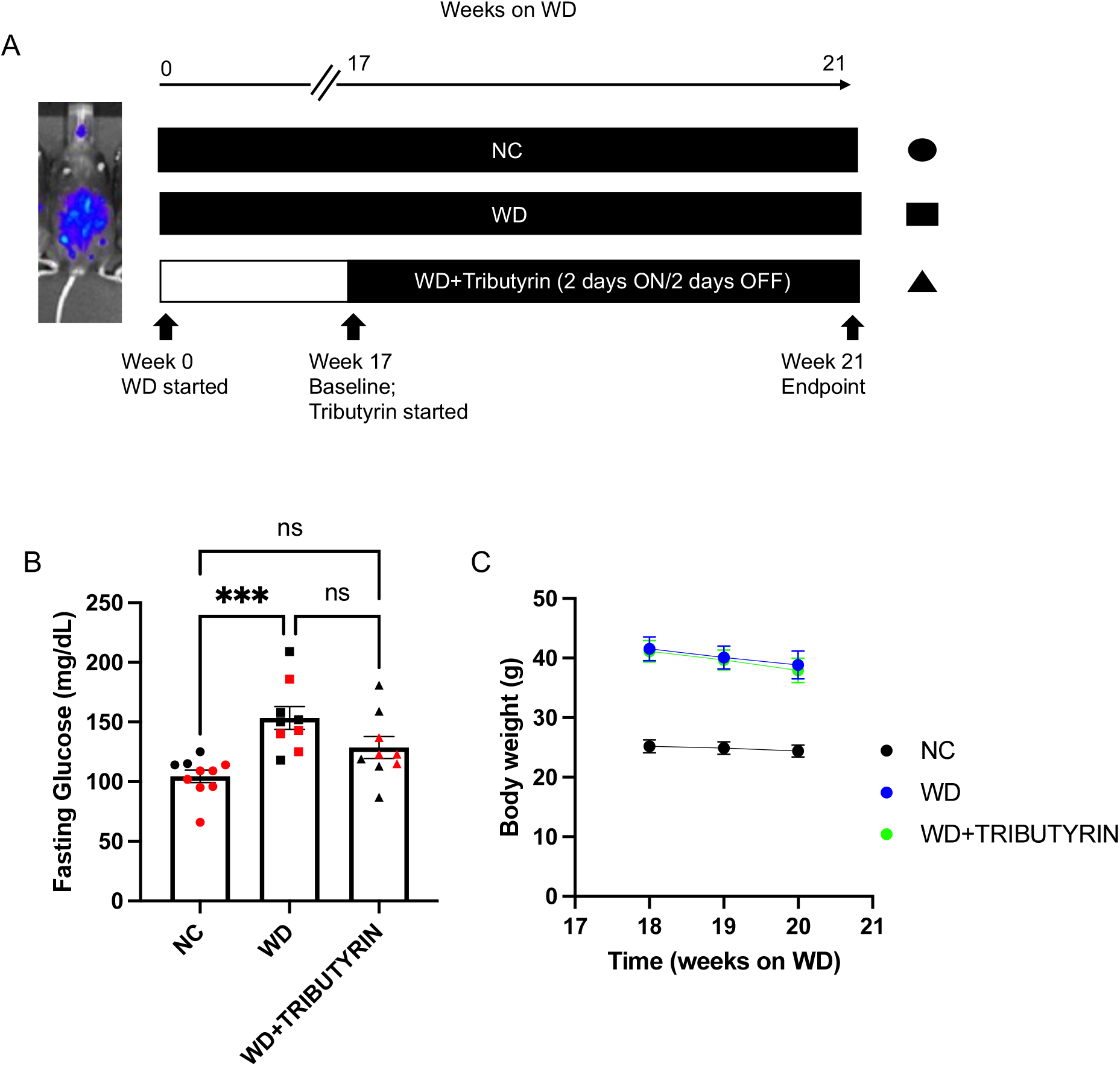
Short Duration of Tributyrin Treatment Trends Towards Improvement of Fasting Glucose Levels. (A) Experimental paradigm. (B) Baseline overnight fasting glucose in NC, Western diet (WD), and WD-fed tributyrin treated male and female mice. (C) Body weight of NC, WD and WD-fed tributyrin-treated mice during the two weeks of tributyrin treatment. One way-anova among NC, WD, and WD+Tributyrin and Tukey’s post-hoc. All values represent mean±S.E.M, *p<0.05, **p<0.01, ***p<0.001, ****p<0.0001, n=4-8 mice/group; males and females.

Next, we investigated the tissue-specific effects of NC, WD and tributyrin on biosensor activation *ex vivo*. In the CNS, the bioluminescence signal was significantly elevated in whole brain of WD-fed male mice when compared to NC and decreased following tributyrin treatment (Figure 4, A–B). We did not detect alterations in bioluminescence signal among the three groups when gating specifically in the hypothalamus and brain stem (Figure 4, D and F), two important areas for neural control of body weight and glucose homeostasis (1, 33). These results suggest that the differences in bioluminescence in whole brain could be due to caspase activation in distinct brain regions, such as the pre-frontal cortex. To complete these observations, we performed flow cytometry studies (gating strategy in supplemental Fig 2) to identify potential changes in immune cell populations in the whole brain of the three experimental groups and did not observe significant differences (not shown). However, we found that the mean intensity fluorescence (MIF) of the pan-leukocyte marker, CD45, was significantly higher in the hypothalamus of obese mice subjected to tributyrin in comparison to WD controls (Figure 4, H), while there were no significant differences in CD45 MIF between NC and WD-fed animals. Because CD45 MIF is indicative of immune cell populations within the brain –CD45^low^ indicating resident microglia and CD45^hi^ indicating infiltrating monocytes (34, 35) – our results suggest a tributyrin-induced recruitment of immune cells in the hypothalamus that was not associated with caspase 1 activation.

**Figure 4.**
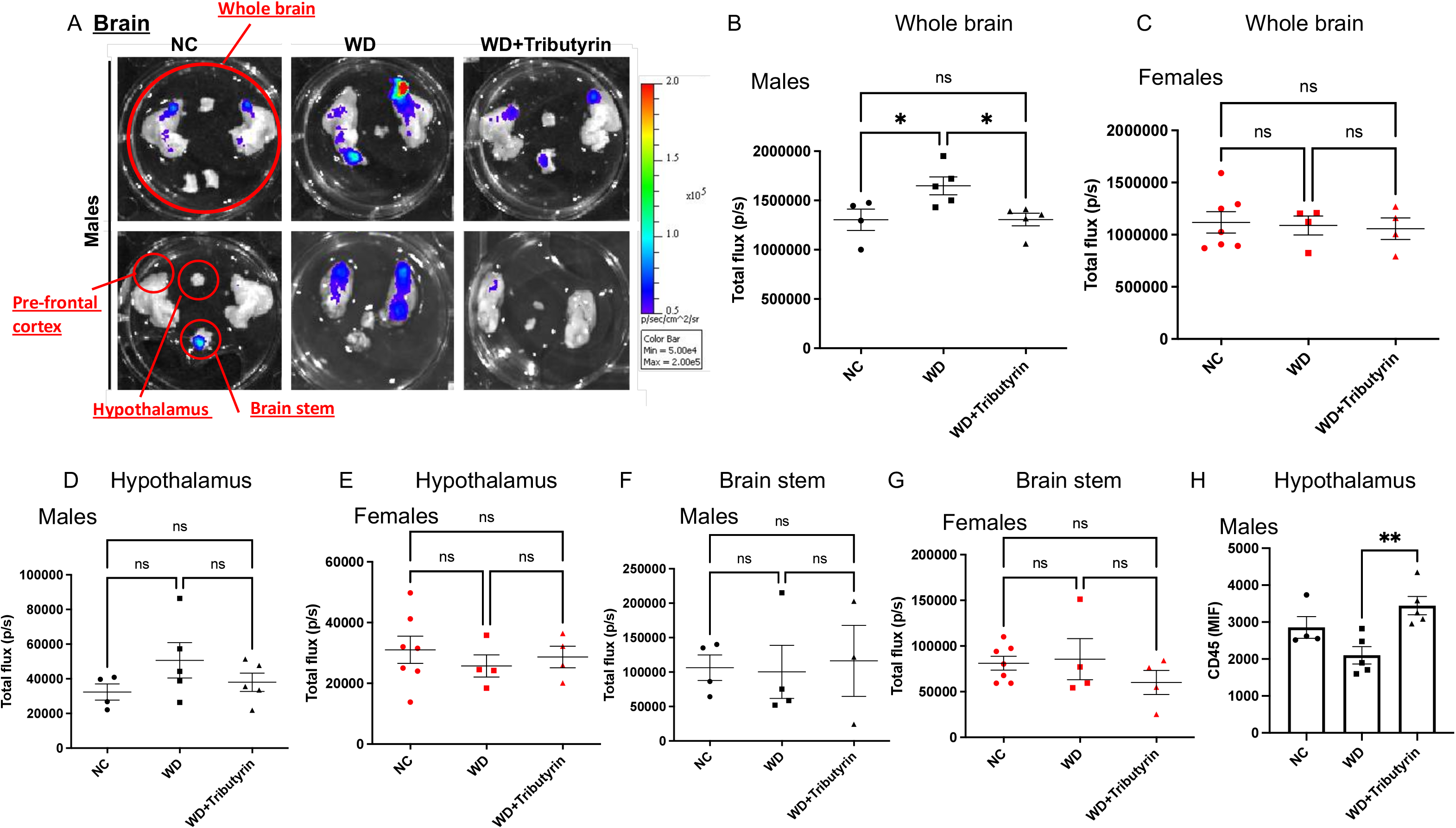
Tributyrin Lowers Caspase-1 Activation and Increases Pan-leukocyte Marker Expression in Whole Brain of Male Mice. (A) Representative IVIS images of brains from normal chow (NC), Western diet (WD) and WD-fed tributyrin treated male mice. (B) and (C) Bioluminescence quantification of whole brain IVIS images for males and females, respectively. (D) and (E) Bioluminescence quantification of hypothalamus IVIS images for males and females, respectively. (F) and (G) Bioluminescence quantification of brain stem IVIS images for males and females, respectively. (H) Mean intensity fluorescence (MIF) of the pan-leukocyte marker CD45 in hypothalamus of NC, WD and WD-fed tributyrin treated male mice. One way-anova among NC, WD, and WD+Tributyrin and Tukey’s post-hoc. All values represent mean±S.E.M, *p<0.05, n=4-8 mice/group; males and females.

We assessed biosensor activation in male and female mice. Interestingly, we did not detect changes in bioluminescence signal among the three groups in females’ whole brain, hypothalamus, and brainstem (Figure 4, C, E, G). Our data reveals that WD and tributyrin modify caspase-1 pathways in a sex-specific manner in the brain of obese mice. Sexual dimorphism has been described in obesity, in which sex hormones play a crucial role in fat depot distribution and function (36). Our data suggest that caspase-1 pathways may underlie some of the sexual dimorphism observed in the obese brain.

The literature describes that butyrate have effect on cardiovascular diseases (19), glucose regulation, thermogenesis, and hepatosteatosis (6, 37, 38). Thus, we aimed at evaluating caspase-1 biosensor activation in metabolic tissues including white and brown adipose tissues (WAT and BAT, respectively), liver, heart, pancreas, and intestines. *In vivo* biosensor activation localized mainly in the adipose tissue surrounding the peritoneal cavity, including subcutaneous, gonadal, and mesenteric fat depots of both, male and female mice (Figure 2A). Thus, we analyzed WAT from NC, WD and WD-fed tributyrin-treated animals and verified a significant increase in total flux of bioluminescence signal between NC and WD in WAT from both, males and females likely due to the increase in total abdominal fat (Figure 5, A, B, and D). However, because WAT size varied among animals belonging to the different experimental groups, we also analyzed the maximum radiance emitted by the tissue (p/s/cm2/sr), which is irrespective of tissue size. We did not detect changes in WAT maximum radiance among the experimental groups in either, males or females (Figure 5, C and E).

**Figure 5.**
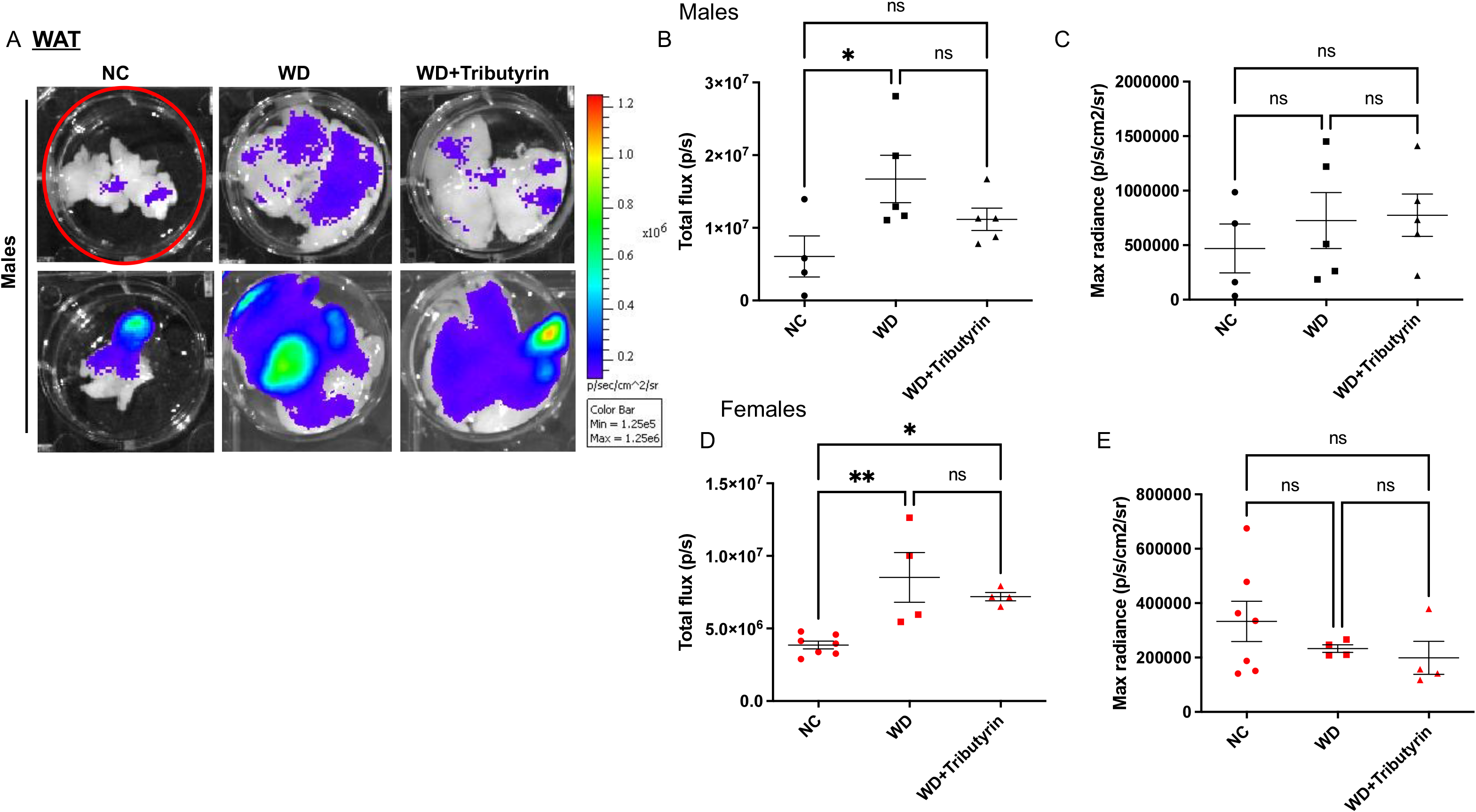
Western Diet Increases Caspase-1 Activation in the White Adipose Tissue (WAT). (A) Representative IVIS images of WAT from normal chow (NC), Western diet (WD) and WD-fed tributyrin treated male mice. (B) and (C) Total flux and maximum radiance bioluminescence quantification of IVIS images for males. (D) and (E) Total flux and maximum radiance bioluminescence quantification of IVIS images for females. One way-anova among NC, WD, and WD+Tributyrin and Tukey’s post-hoc. All values represent mean±S.E.M, *p<0.05, **p<0.01, ***p<0.001, n=4-8 mice/group; males and females.

However, we observed a significant increase in BAT bioluminescence in WD-fed tributyrin treated male mice compared with NC and WD-fed animals (Figure 6, A–C). The biosensor differences observed in BAT were sex-specific, and not recapitulated in the females (Figure 6, D and E).

**Figure 6.**
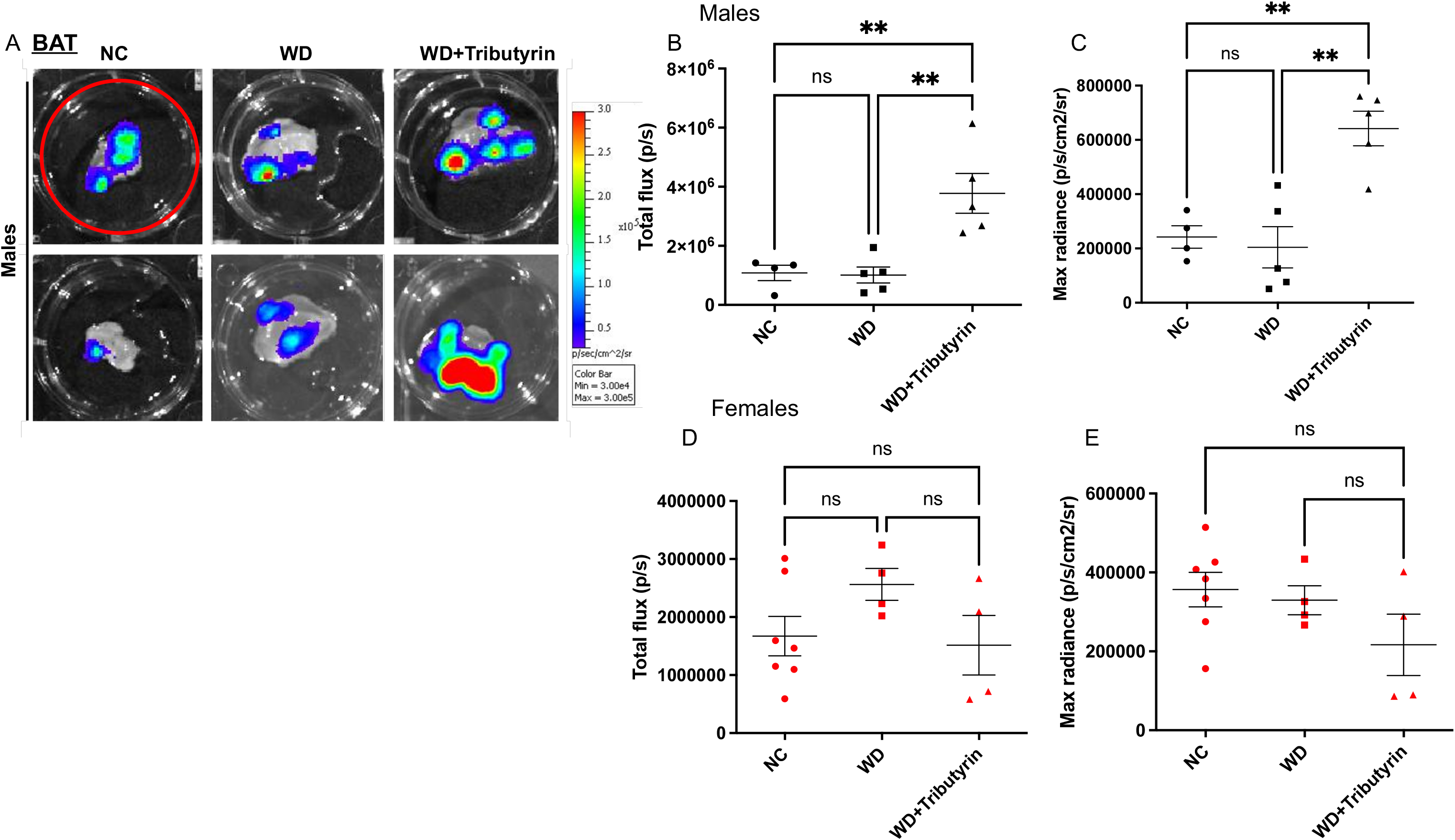
Tributyrin Increases Caspase-1 Activation in the Brown Adipose Tissue (BAT) of Male Mice. (A) Representative IVIS images of BAT from normal chow (NC), Western diet (WD) and WD-fed tributyrin treated male mice. (B) and (C) Total flux and maximum radiance bioluminescence quantification of IVIS images for males. (D) and (E) Total flux and maximum radiance bioluminescence quantification of IVIS images for females. One way-anova among NC, WD, and WD+Tributyrin and Tukey’s post-hoc. All values represent mean±S.E.M, *p<0.05, **p<0.01, ***p<0.001, n=4-8 mice/group;males and females.

We then analyzed biosensor activation in livers of NC, WD and WD-fed tributyrin-treated male and female mice. We detected a significant increase in total flux in liver of obese male mice compared to lean controls, that was unaltered by tributyrin treatment (Figure 7, A and B). There were no significant differences in bioluminescence signal in liver of female animals (Figure 7, C). We also did not detect significant changes in bioluminescence signal from pancreas and intestine in either, males or females (Supplement Fig. 3).

**Figure 7.**
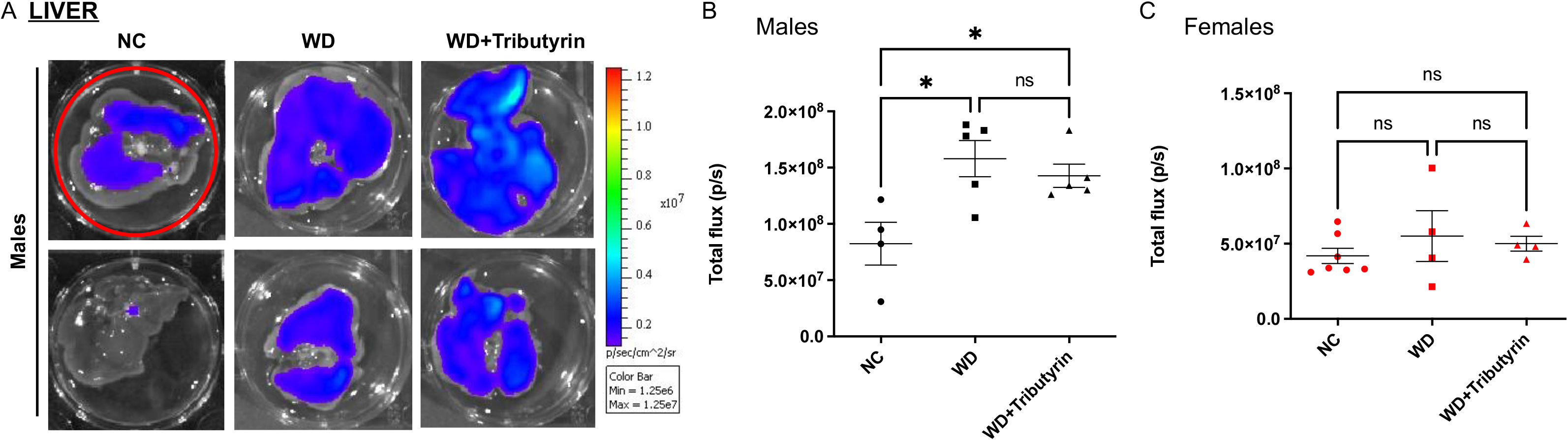
Western Diet Increases Caspase-1 Activation in the Liver of Male Mice. (A) Representative IVIS images of liver from normal chow (NC), Western diet (WD) and WD-fed tributyrin treated male mice. (B) and (C) Total flux bioluminescence quantification of IVIS images for males and females, respectively. One way-anova among NC, WD, and WD+Tributyrin and Tukey’s post-hoc. All values represent mean±S.E.M, *p<0.05, **p<0.01, ***p<0.001, n=4-8 mice/group; males and females.

We measured caspase-1 activation in the heart of animals from the aforementioned groups. We detected changes in caspase-1 activation in both males and females (Figure 8, A and B). In both sexes, there was a significant increase in total flux between NC- and WD-fed mice that was attenuated by tributyrin treatment (Figure 8, B). To validate our data, we performed western blot analysis in heart from NC, WD, and WD-fed tributyrin-treated female mice. In agreement with the biosensor data, we measured increased activation of caspase-1 in heart tissue from obese female mice when compared to lean controls, as observed by a significant increase in cleaved caspase-1 amounts (Figure 8, C–D). WD-fed tributyrin-treated females had significant lower levels of cleaved caspase-1 than hearts from WD-fed mice (Figure 8. C–D), indicating an attenuation of caspase-1 activation by tributyrin. We also measured an increased trend in IL-1β production in hearts from WD-fed mice in comparison to lean controls and tributyrin-treated mice (Figure 8, E). These data confirm that biosensor activation was occurring in sites of ongoing inflammasome/caspase-1 activation in this tissue.

**Figure 8.**
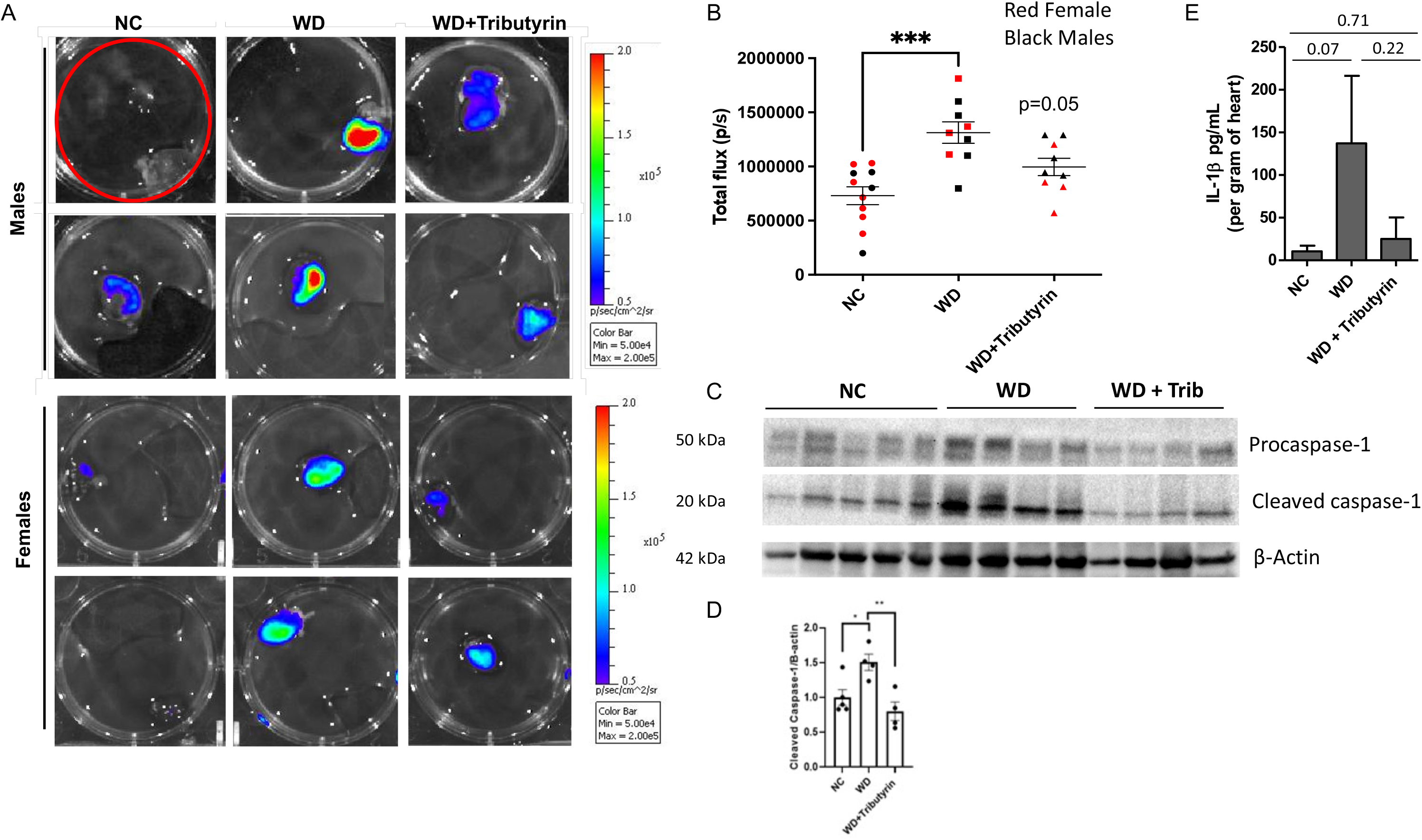
Tributyrin Decreases Caspase-1 Activation in the Heart. (A) Representative IVIS images of heart from normal chow (NC), Western diet (WD) and WD-fed tributyrin treated male mice. (B) Bioluminescence quantification of IVIS images. (C) Immunoblot images of procaspase-1, cleaved caspase-1 and β-actin proteins in heart samples of NC, WD, and WD-fed tributyrin-treated female mice. (D) Densitometric analysis of the ratio of cleaved caspase-1/β-actin protein bands. (E) IL-1 levels in hearts from NC, WD, and WD+Tributyrin female mice. One way-anova among NC, WD, and WD+Tributyrin and Tukey’s post-hoc. All values represent mean±S.E.M, *p<0.05, **p<0.005; n=4-8 mice/group; males and females.

### Butyrate stimulation modifies expression of proinflammatory markers in bone marrow-derived macrophages from obese mice

We hypothesized that the changes in caspase-1 observed in WD-fed mice and after tributyrin treatment may be due to an effect of butyrate on macrophages. Thus, we stimulated bone marrow-derived macrophages (BMDM) from WD-fed mice with lipopolysaccharide (LPS) –in the presence or absence of sodium butyrate (NaB). Exposure to NaB significantly decreased LPS-induced mRNA expression of pro-caspase-1, IL-6 and TNFa (Figure 9, A, C, D) as well as IL-6 secretion (Fig. 9E). While pro-IL-1β mRNA expression was significantly higher in LPS-stimulated BMDMs compared to non-treated macrophages, NaB exposure did not reduce expression of this proinflammatory transcript (Figure 9, B). Collectively, these results suggest that butyrate may have a protective role in preventing LPS-induced activation of inflammatory responses in macrophages in obese mice, partially by decreasing expression and secretion of some inflammatory cytokines.

**Figure 9.**
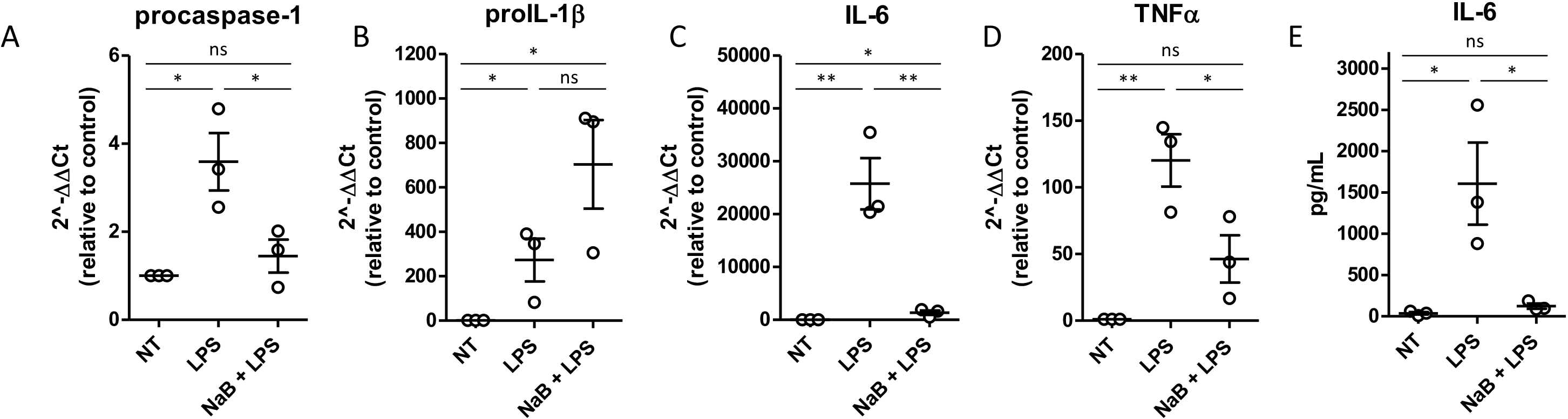
Butyrate Modifies Cytokine Expression in Bone Marrow-Derived Macrophages (BMDM). (A) Pro-caspase-1, (B) pro-IL-1β, (C) IL-6, and (D) TNF-a mRNA levels in BMDM from Western diet (WD)-fed mice following lipopolysaccharide (LPS) stimulation or LPS/sodium butyrate (NaB) co-exposure. (E) IL-6 levels in media from BMDM from WD-fed mice following LPS or LPS+NaB treatment; One way-anova and Tukey’s post-hoc. All values represent mean±S.E.M, *p<0.05, **p<0.01, ***p<0.001, ****p<0.0001, n=3 independent experiments; 3 mice/group; males.

## Discussion

Herein, we investigate the effects of WD feeding and butyrate treatment on whole body and tissue-specific inflammation. The method employed allowed for monitoring of caspase-1-mediated inflammation longitudinally in vivo. Thus, the presented work suggests that WD feeding leads to a gradual increase in caspase-1 activation in vivo over time. The bioluminescence signal originated from activated luciferase/caspase-1 construct allowed for visualization and quantification of inflammation in the animals and it was mainly coming from in the abdominal cavity. This result implies that increases in inflammation in obese mice could be occurring in: (i) the white adipose tissues – subcutaneous and visceral fat –, (ii) internal organs, including pancreas and intestines, or (iii) the peritoneal cavity – loaded with immune cells, such as macrophages. Thus, to confirm the sites of increased caspase-1 activation, we analyzed tissues ex vivo at the study endpoint. Remarkably, we detected differences in bioluminescent signal in the fat tissue, as well as in other metabolic tissues, that could be responsible for the increase in inflammation observed in vivo. Ventrally, there were significant changes among the experimental groups in the liver and heart; there were no differences in the remaining anteriorly located organs analyzed (the brown adipose tissue was collected from the posterior cervical region). Therefore, these data indicate that the bioluminescence signal is likely originating in cells within the peritoneal cavity – a major hub of circulating immune cells – overlooked in our studies.

Butyrate effects observed in the brain can be explained by the gut-brain axis – the bidirectional communication between the CNS and the gastrointestinal system (18, 39, 40). Through vagal afferents and by crossing the blood brain barrier via monocarboxylate transporters (MCT) expressed on endothelial cells, butyrate can exert specific functions in certain brain nuclei, such as the frontal cortex and the hippocampus (18, 39, 40). For instance, butyrate can act as endogenous ligand for free fatty-acids receptors (FFARs – G-coupled protein receptors) or it can also act as a histone deacetylase inhibitor (HDACi) and enhance cell-specific gene transcription by opening chromatin conformation (18, 39, 40). The whole-brain biosensor data revealed a significant increase in caspase-1 mediated inflammation upon WD feeding that was significantly attenuated by tributyrin treatment. Although it is known that WD can severely impact brain function, including memory and cognition, we were able to visually suggest caspase activation following 20 weeks of WD feeding paradigm. At that time point, the systemic low-grade inflammation seen in obesity likely induced tissue-specific inflammatory response, including within the CNS. Flow cytometry analysis of the hypothalamic region also revealed a distinct immune cell profile influenced by diet and tributyrin. The fluorescence levels of the pan-leukocyte marker, CD45, can be indicative of specific immune cells within brain regions i.e., lowly expressing CD45 cells (CD45^low^) represent resident microglia, while highly expressing CD45 cells (CD45^hi^) characterize infiltrating blood-derived monocytes, such as perivascular and meningeal macrophages (34, 35). NC- and WD-fed animals had low CD45 fluorescence, suggesting the presence of resident microglia population (34, 35). However, obese mice subjected to tributyrin treatment presented significantly higher CD45 fluorescence levels in comparison to WD controls, which is characteristic of perivascular and meningeal macrophages (34, 35). These differences in immune cell profile could in turn suggest the overall level of inflammatory response within the hypothalamus. The hypothalamus is a region of particular interest because of its fundamental role in host metabolism and energy balance, controlling metabolic functions – such as sleep, thermoregulation, and hunger – through the secretion of neuroendocrine molecules (33). Within the hypothalamus, nuclei in close proximity to the third ventricle, for instance the arcuate nucleus, are most likely to suffer the effects of circulating compounds, including butyrate, due to a less restricted blood-brain barrier (33). Perivascular and meningeal macrophages are blood-derived monocytes that infiltrate neural tissues, while microglia are the CNS resident glial cells. Perivascular macrophages recruitment to the hypothalamus can be part of an adaptive immune response to systemic inflammation, since they have been implicated in activation or suppression of the HPA axis in response to inflammatory stimuli (34, 35). Thus, our results suggest that the differential effect of tributyrin could be driven by particular immune cell populations residing/infiltrating these organs. Centrally, tributyrin might exert anti-inflammatory actions by potentially decreasing caspase-1 activation in the brain of obese mice and by potentially increasing perivascular and meningeal macrophages (CD45^hi^) in the hypothalamus. Differently, tributyrin does not alter caspase-1 activation in the PNS but it does act on macrophage infiltration and polarization.

The main function of BAT is thermogenesis – to produce heat by dissipating energy. Uncoupling proteins (UCPs) are the drivers of BAT thermogenesis, as they dissipate mitochondrial proton gradient and allow the uncoupling between the respiratory chain and ATP generation (41). However, thermogenesis was impaired in mice fed high-fat diet (58% fat) due to BAT inflammation and damage involving ER stress and mitochondrial dysregulation (42). In our WD model (42% fat), we did not observe changes in biosensor signal in BAT of WD- compared to NC-fed mice. We did, however, see a significant increase in caspase-1 activation in BAT of obese male mice following tributyrin treatment. In this case, tributyrin seems to enhance inflammatory response by activating caspase-1 signaling. Li et al. demonstrated that oral butyrate treatment led to increased BAT thermogenic capacity (37). In that work, the authors showed that chronic butyrate administration activated BAT by increasing UCP-1 mRNA expression and via increased sympathetic outflow, leading to improvements of some high-fat diet phenotype, such as glucose metabolism and insulin signaling (37). Therefore, the changes in inflammatory response observed with our model likely precede the behavioral changes that would occur with a longer tributyrin paradigm.

The biosensor data obtained from hearts were the only results recapitulated in female mice. In both sexes, we observed a significant increase in caspase-1 activation upon 20 weeks of WD that was significantly attenuated following two weeks of tributyrin administration. Obese individuals are at higher risks for developing cardiovascular disease and the systemic low-grade inflammation is known to impair cardiac remodeling (43). Immune cell infiltration and M1 macrophage polarization, both associated with obesity and Metabolic Syndrome (MetS) components, lead to cardiac injury. (43) In addition, gut microbiome alteration in obesity has recently been linked to cardiovascular disease and myocardial infarct (44). Obese mice transplanted with cecal contents from lean animals presented reduced myocardial infarct size, concomitant with alterations in gut permeability and cecal butyrate content (44). Thus, our findings, in agreement with the literature, indicate that not only does WD lead to increased caspase-1 mediated inflammation of the cardiac tissue, but also that tributyrin is able to rescue WD-induced cardiac injury by reducing inflammation within the heart. Increased active caspase-1 and IL-1β levels in the hearts of WD-fed mice confirms that there is ongoing inflammasome activation in this tissue. Further experiments to identify the inflammasome(s) driving these responses in the heart during obesity are needed.

NaB exposure decreased the expression of proinflammatory transcripts, pro-caspase-1, IL-6 and TNFa in LPS-stimulated BMDMs isolated from WD-fed mice. We also detected reduced secretion of IL-6. We can infer that butyrate seems to confer protection against LPS-induced pro-inflammatory response – and possibly against other insults.

Although a powerful technique to assess inflammation dynamics *in vivo* and tissue-specific capsase-1 mediated inflammation, a drawback of our method is the lack of cell specificity. Thus, in future studies we aim to assess what cell type (s) are indeed involved in caspase-1 mediated inflammation within the tissues studied, providing a further knowledge on WD-induced inflammation and butyrate action.

Previous studies have reported on the sexual dimorphism of obesity (36). Differential adipose tissue distribution and function between men and women that lead to distinct metabolic and neural features are mainly driven by the hormonal patterns in both sexes (36). Some of the results presented highlight WD-associated sex differences regarding caspase-1 activation. Most of the tissue-specific analyses revealed alterations in inflammation in male only in addition to different time-course. These data agree with the literature highlighting that female are more resistant to WD than males, but they develop earlier complications such as insulin resistance likely associated with inflammation (45). This suggests a differential regulation of caspase-1 inflammation between males and females upon WD feeding and butyrate treatment, potentially driven by sex hormones.

Given our hypothesis that changes in caspase-1 activity due to butyrate were mediated by macrophages, we investigated putative pro-inflammatory pathways that might be suppressed by the short-chain fatty acid. Toward this goal, we performed a computational analysis of a manually-curated signaling network for macrophages. Our procedure consisted of four steps: 1) identification of LPS-mediated pathways culminating in caspase activity, 2) a query for genes susceptible to butyrate, 3) identifying butyrate-mediated genes that are linked to the LPS-mediated pathways using protein-protein interaction pairs (PPIs) identified from PPI database and 4) traversing the pathways to identify those suppressing caspase-1 activity. Details of this approach, including the database and the network traversal algorithm, are described in Supplemental Methods.

Our curated network reflects that classical M1 activation via LPS/TLR4 complex promotes the activation of NF!B p65/p50 (associated gene, NFKB1), which initiates gene transcription associated with pro-inflammatory cytokines or inflammatory species, such as IL-1β or NLRP3/caspase-1 complex (46, 47), respectively. Our pathway analysis based on these databases suggests that the FFAR2-mediated pathway, when stimulated by butyrate, suppresses the LPS-mediated classical activation of macrophages. This occurs through butyrate stimulation of the G protein-coupled receptor (GPCR) FFAR2 (48), which promotes beta-arrestin 2 (associated gene, ARRB2) activity and ultimately inhibition of the NF!B complex. ARRB2 has previously been established to suppress NF*κ*B activation via promoting its inhibition by NF!B inhibitor alpha (associated gene, NFKBIA) (49, 50). Hence, reduction of caspase-1 in tributyrate-treated WD-mice may stem from FFAR2-mediated suppression of NFkB activity. The analysis also indicated that butyrate may directly inhibit HDAC3 activity (51), which also results in suppressing the transcription of pro-inflammatory proteins via NF*κ*B activation.

Our work also demonstrated that WD feeding lead to a gradual increase in caspase-1 activation in vivo over time. Although this work only assessed changes in biosensor signal following a short therapeutic intervention of tributyrin in WD-fed mice, future work should measure the long-term effects of tributyrin administration on the development and/or resolution of caspase-1-dependent inflammatory responses *in vivo* and in specific tissues *ex vivo*. Thus, our study unmasked that the biosensor mice can be extremely useful to perform tissue specific drug discovery that would improve complications of obesity impacting specific tissue.

### Animal studies

Animal studies were conducted in accordance with recommendations in the Guide for the Care and Use of Laboratory Animals of the National Institutes of Health and with the approval of the Loyola University Chicago Institutional Animal Care and Use Committee. Wild type, male C57BL/6J (#000664) mice were obtained from Jackson Laboratories (Maine, USA). Caspase-1 biosensor mice were generated in-house as described elsewhere (1). Animals were housed under a 12:12 hour light/dark cycle. The lean group was fed Normal Chow (NC) (Teklad LM-485), while the experimental groups were fed Western Diet (WD) (TD88137, Teklad Diets; 42%kcal from fat, 34% sucrose by weight, and 0.2% cholesterol total - Envigo, Indiana, USA) for 20 weeks starting at 13 weeks of age.

### Tributyrin treatment

13-week old mice were fed NC or WD for 20 weeks. At the end of week 17, animals were needle fed with tributyrin (3 butyrate molecules attached to a glycerol backbone. 5g/kg body weight. Sigma-Aldrich)) or 1X PBS (used as vehicle) every 48 hours (2 days on/2 days off) for 2 weeks (n=4-8 mice per group).

### Fasting glucose

Mice were overnight fasted and glucose was measured from tail blood drops using AlphaTrak glucometer for rodents (Fisher Scientific, Pennsylvania, USA).

### IVIS imaging

In vivo imaging was done as previously described (1). Briefly, mice were anesthetized, weighed and injected i.p. with a single dose of 150 mg/kg VivoGlo Luciferin. Anesthetized mice were imaged with the IVIS 100 Imaging System (Xenogen) 10 min after administration of the luciferase substrate. For ex vivo imaging, mice were sacrificed immediately following in vivo image acquisition and dissected tissues were placed in a 6-well plate with 1mL of luciferase substrate (300μg/mL). Bioluminescence was quantified and analyzed use Living Image software.

### Flow cytometry

Following euthanasia, hypothalamus, DRG and SN were dissected and placed in trypsin/collagenase A (1.25mg/ml each – Sigma Aldrich) in 1X Phosphate-buffered saline (PBS) and incubated for 30 min at 37°C in a 5%CO_2_ incubator. Enzymatically digested tissues were carefully resuspended by the use of glass pipettes and dissociated cells were pelleted by a 5 min centrifugation at 300x *g*. The supernatant was carefully removed and cells were placed in blocking solution (5% horse serum in 1X PBS) for 30 min on ice. After pelleting the cells at 300x *g* for 5 min, fluorescently conjugated antibodies were added at 1:1000 dilution (anti-CD45, anti-CD11b, anti-F4/80, anti CD206, anti-IA-IE – Biolegend) and incubated for 30 min on ice. Cells were then washed and resuspended in 5% horse serum 1X PBS. Flow cytometry data was acquired using BD FACS Aria III and data was analyzed using FlowJo (Treestar).

### Western Blot Analysis

Protein from frozen mouse hearts were prepared by lysis in ice-cold RIPA buffer (ThermoFisher, cat.89900) containing protease and phosphatase inhibitor (ThermoFisher, cat. A32959). Tissue was homogenized using bullet blender bead lysis kit (Next Advance), and protein concentrations were determined with Pierce BCA Protein Assay Kit (Thermofisher, cat. 23225). Proteins were separated by sodium dodecyl sulfate polyacrylamide gel electrophoresis (SDS-PAGE) on 4-15% gradient gel (Bio-Rad, cat.4561086) and transferred to PVDF membrane using iBlot 2 transfer system (ThermoFisher). Protein expression was measured by chemiluminescence using ChemiDoc imaging system (Bio-Rad). Proteins were detected with the following primary antibodies: anti-Caspase-1(p20) (AdipoGen), anti-B-actin (8226, abcam).

### Culture and treatment of murine bone marrow-derived macrophages (BMDM)

Bone marrow cells were isolated from the femurs and tibias of WD-fed mice and cultured in medium containing 20 ng/mL M-CSF, as previously described (Talley paper, and PMID 29177864). After 6 days in culture, differentiated bone marrow-derived macrophages (BMDMs) were re-plated in 24 well plates and treated with 100ng/mL LPS (from *Escherichia coli* O26:B6; Sigma-Aldrich) with or without 1mM sodium butyrate for 4 hours.

### Cytometric Bead Array (CBA)

For cytokine analysis in tissues, tissues were processed as described above and homogenates were centrifuged and lysate collected. For in vitro experiments, supernatant was collected from BMDMs following treatments. CBA was performed using BD CBA Flex sets for the indicated cytokines (BD Biosciences) following the manufacturer’s instructions and data were analyzed on an LSRFortessa (BD Biosciences).

### Quantitative PCR

Following treatment, BMDMs were washed once with 1X PBS (Corning), and RNA was isolated using the RNEasy Mini Kit (Qiagen). To obtain cDNA, a reverse transcription reaction was performed using Taq polymerase reverse transcription kit (Applied Biosciences). For all genes of interest (listed in Table 5), qPCR was performed using Sybr green-based assay (Roche, Indiana, USA) using IDT primers (IDT technologies, Iowa, USA). Actin was used to normalize data and quantification was done using ΔΔCT method with vehicle treated group’s mean value set at 100% as reported before (2).

**Table 5.**
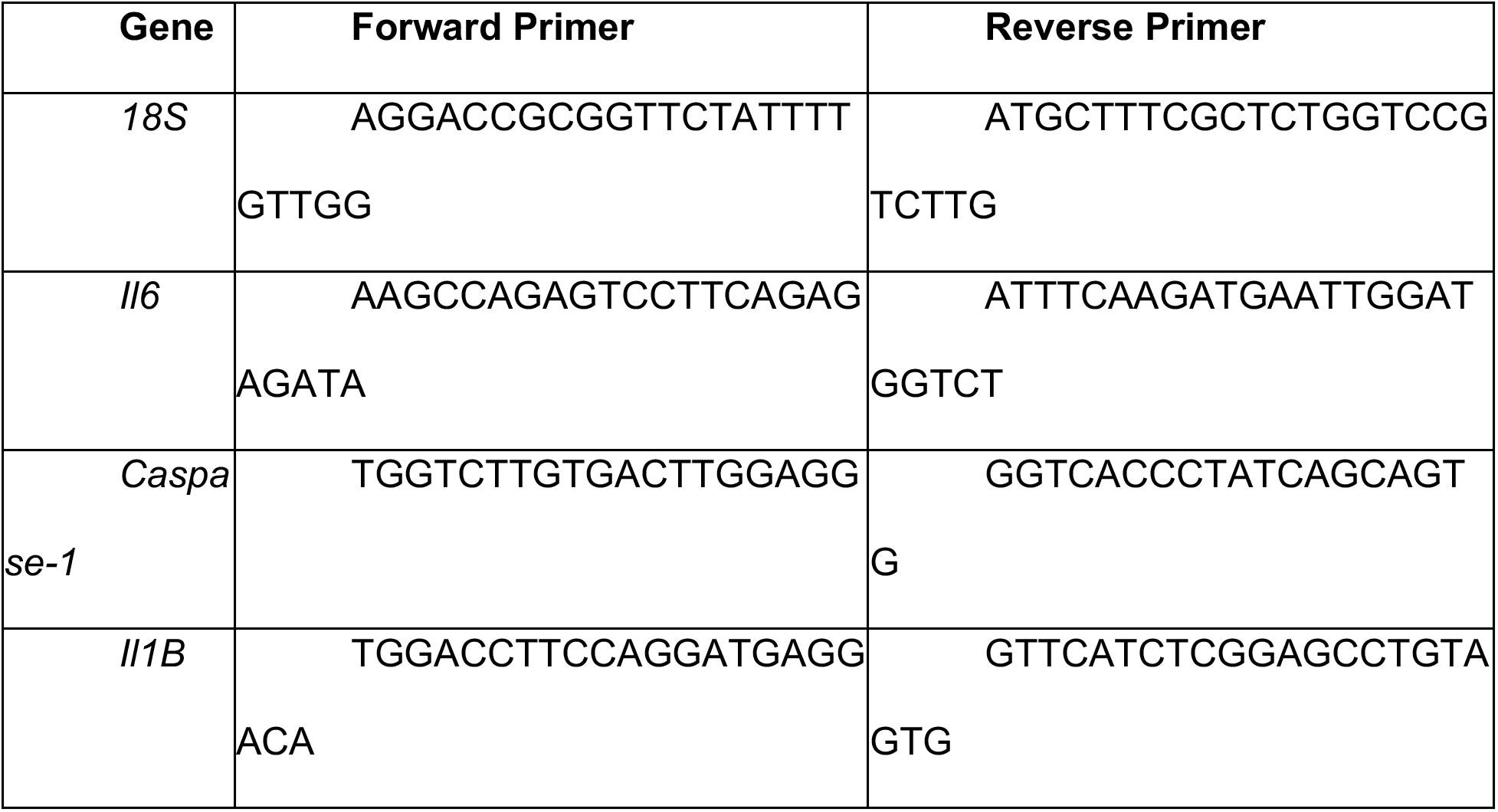
Primer Sequences for Genes of Interest.

### Pathway analysis

The manually-curated network was built from several databases including KEGG, STRING, and SIGNOR (3–5). Kegg was used for identifying LPS-mediated inflammatory networks [7]. The SIGNOR database provides pathways with general classifications, such as “macrophage polarization” or “toll like receptor pathway” (4). The STRING database provides sets of protein-protein interactions collected from the scientific literature, as well as genes that are annotated as targets for butyrate (3). Cytoscape (6) was used to merge networks from KEGG and SIGNOR. The STRING database provides a list of genes interacting with the target gene in the descending order that is from the most probable interaction based on its methodology. The PPIs from STRING were added as edges to the graph using NetworkX. Each PPI was designated as activating or inhibitory based on KEGG annotations or review of relevant literature. The resulting graph (see SI Appendix Fig. 4) was subjected to a minimum path analysis via NetworkX to identify pathways linking each butyrate-sensitive gene to caspase-1. Each pathway was marked as activating or inhibitory based on the number of inhibitory PPIs within a path (e.g. an odd number of inhibitory PPIs was deemed as an inhibitory path, otherwise an even or zero number was designated as activating). The resulting inhibitory paths are shown in Fig 10 A Jupyter notebook and supporting python files, as well as the curated network, are available at https://colab.research.google.com/drive/1l_8SHvtZQo1U7SQg_wfUPvwx0_S-drjZ?usp=sharing

**Figure 10.**
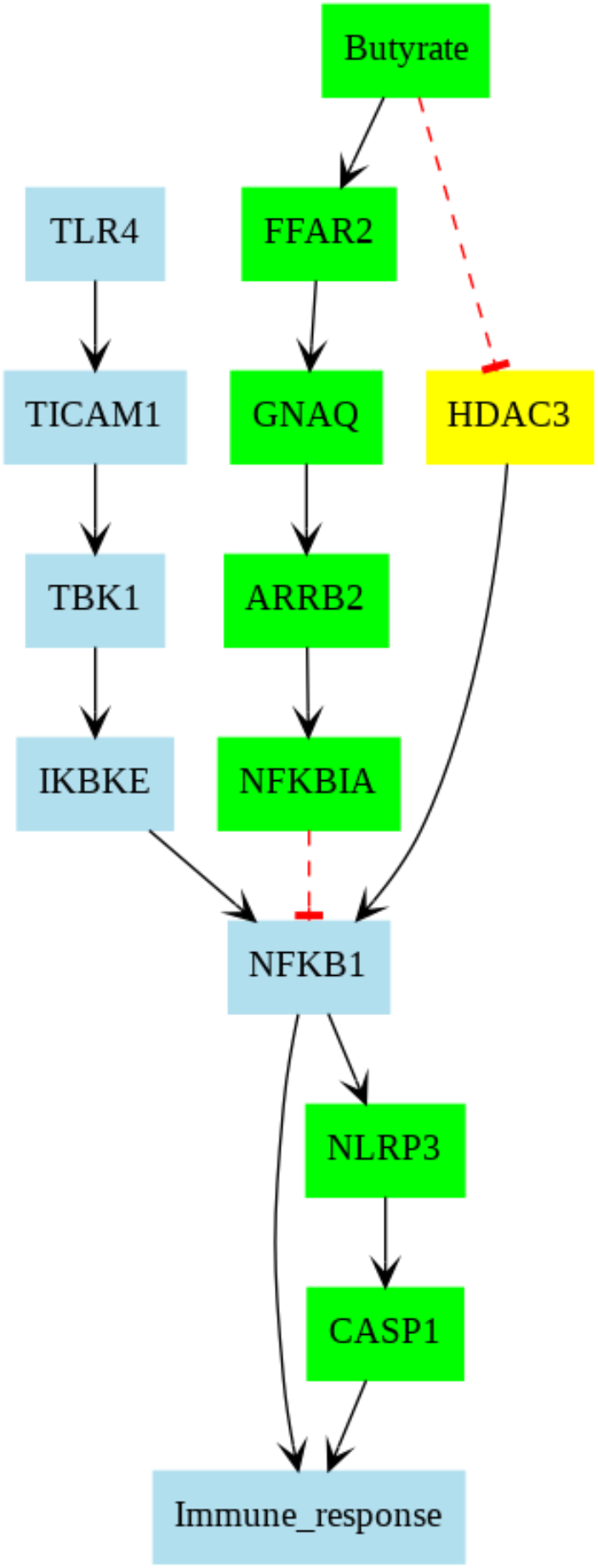
Pathway collected from STRING database. Each color is contributed by a specific pathway. Black and red-colored pathways are from LPS/TLR4- and FFAR2-mediated pathways, respectively. The black arrow indicates the promoting correlation whereas the red line with the tee marker denotes the inhibitory correlation

### Statistical analysis

All data are represented as mean ± S.E.M. Analyses were performed using Graphpad Prism. For single group comparisons a 2-tailed t-test or Mann-Whitney test were used as appropriate and multiple comparisons were performed using ANOVA between groups with differences identified by post-hoc tests as shown in corresponding figure legends. p< 0.05 was considered significant.

## Acknowledgments

The authors would like to acknowledge Patricia Simms from Loyola University FACS Core. This work was supported by 1R01DK117404-01 and DIACOMB Pilot & Feasibility project to V.M.A.

## Competing Interest Statement

The authors declare no competing interest.

## Supplement Figure Legends

**Figure 1.**
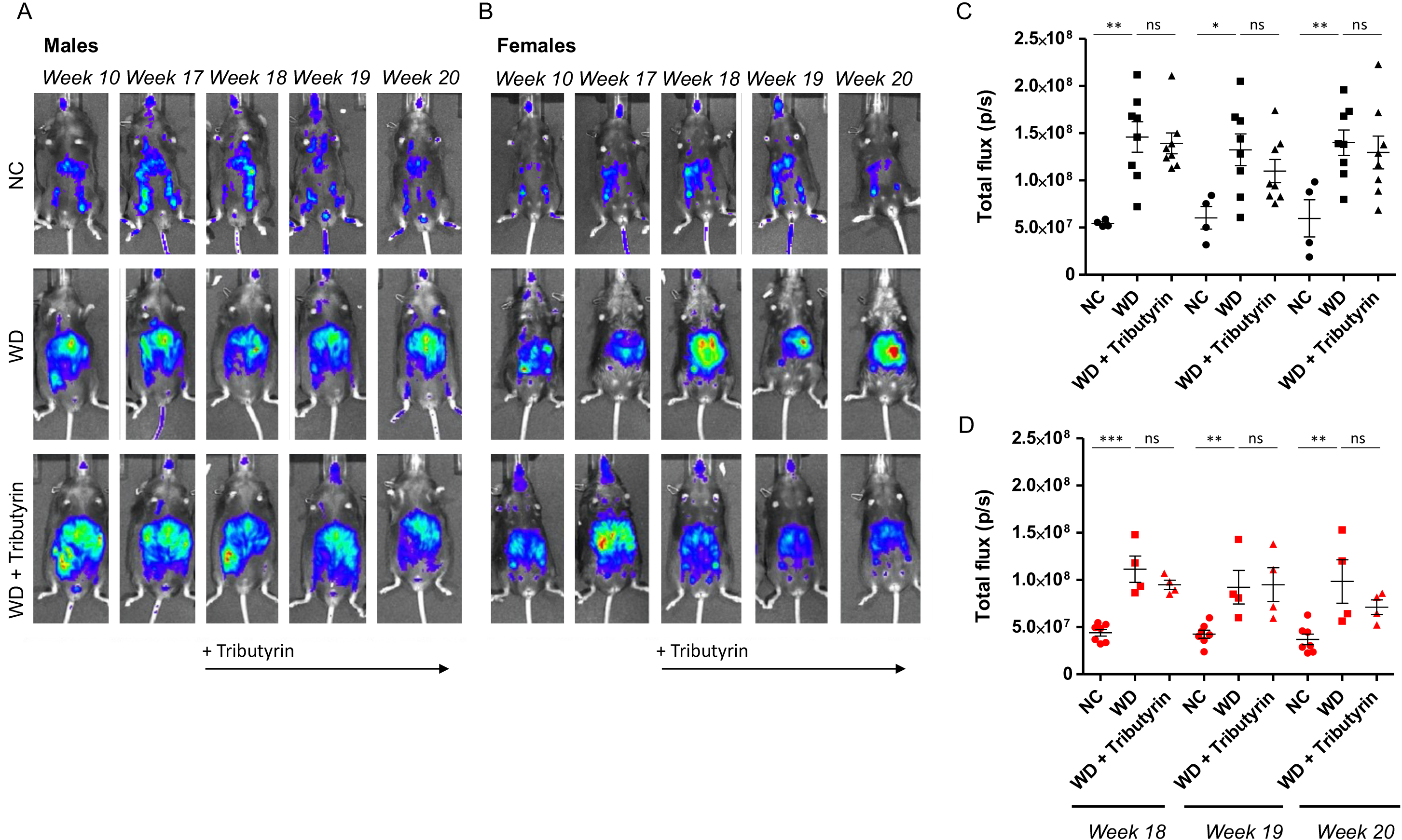
Tributyrin Does not Affect Overall Caspase-1 Activation In Vivo. (A) and (B) Representative whole body IVIS images from normal chow (NC), Western diet (WD) and WD-fed tributyrin-treated male and female mice over time. (C) and (D) Bioluminescence quantification of respective IVIS images. One way-anova among NC, WD, and WD+Tributyrin and Tukey’s post-hoc for every timepoint. All values represent mean±S.E.M, *p<0.05, **p<0.01, ***p<0.001, n=4-8 mice/group; males and females.

**Figure 2.**
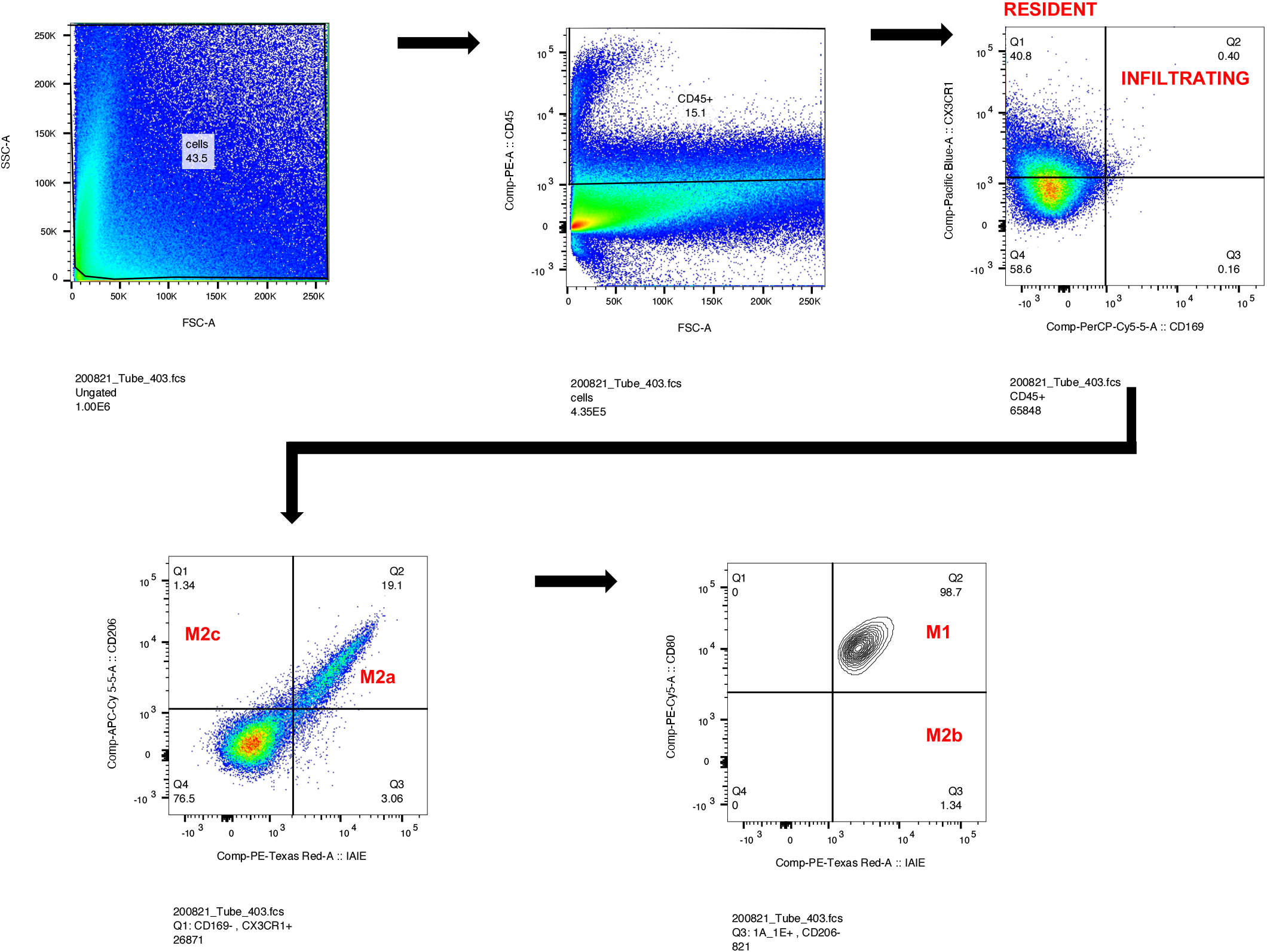
Microglia Gating Strategy. Flow cytometry gating strategy for tissues in the central nervous system.

**Figure 3.**
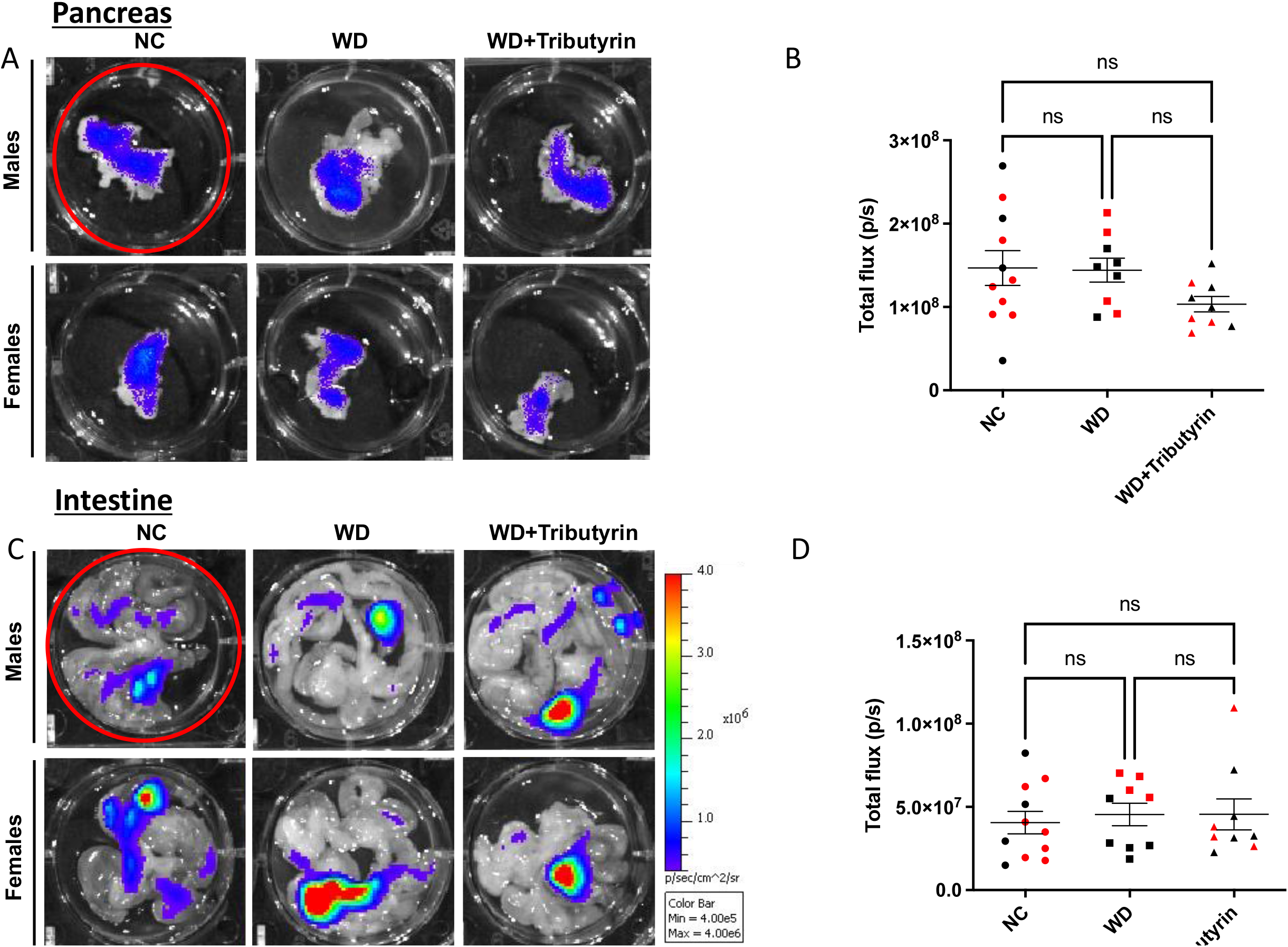
Tributyrin Treatment Does Not Alter Caspase-1 Activation in Pancreas, and Intestines of Male and Female Mice. IVIS quantification of (A) pancreas and (C) intestines of normal chow (NC), Western diet (WD) and WD-fed tributyrin-treated male and female mice. (B) and (D) Bioluminescence quantification of pancreas and intestine IVIS images, respectively. All values represent mean±S.E.M, *p<0.05; n=4-8 mice/group; males and females.

**Figure 4.**
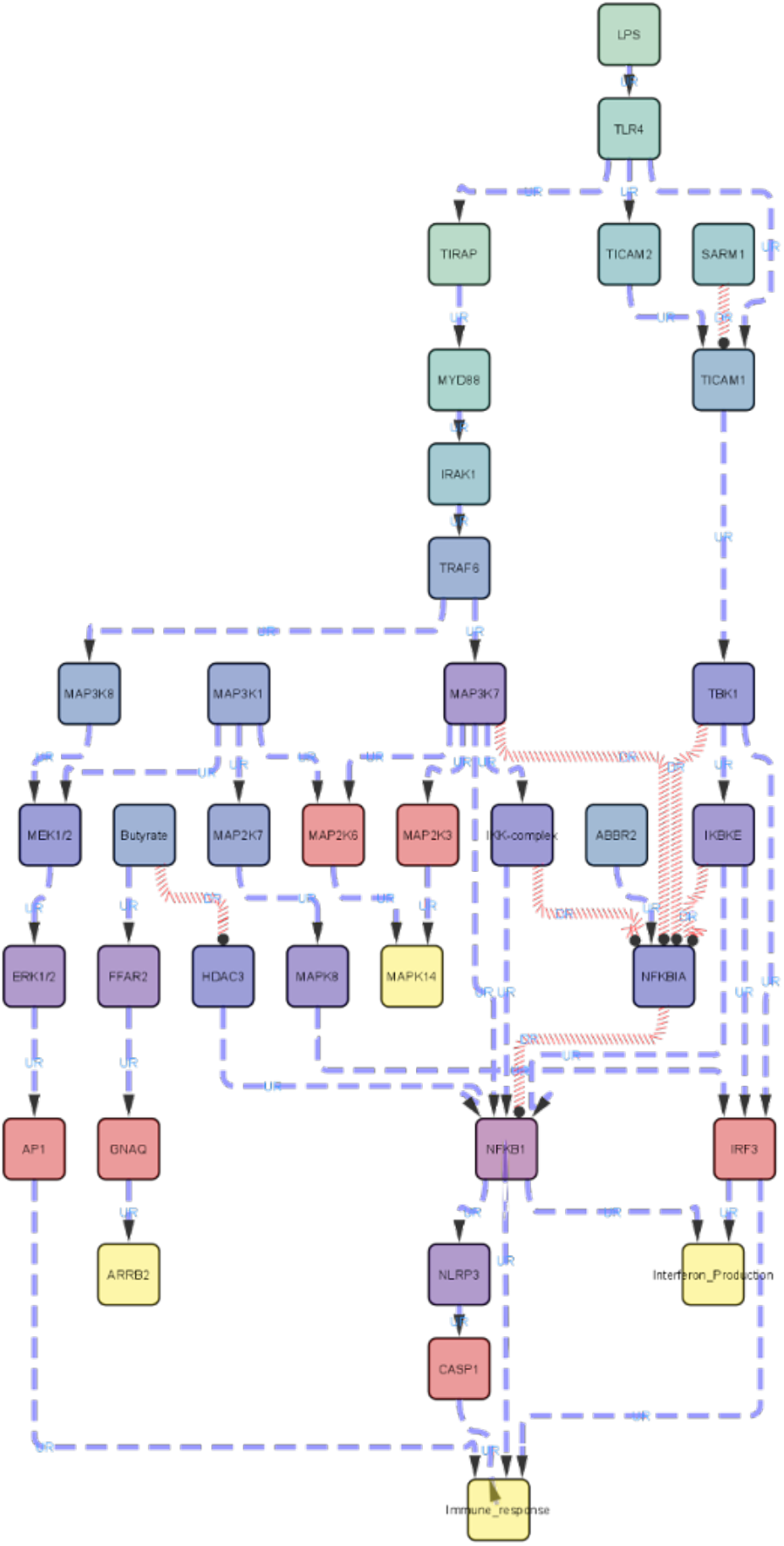
LPS/TLR4 complex-mediated pathway collected from KEGG, SIGNOR, and STRING and visualized by Cytoscape. FFAR2 and HDAC3 pathways were added via extended search in STRING database. Blue colored edges (path) indicate promoting whereas red colored edges indicate inhibitory interaction.

## Notes

### Competing Interest Statement

The authors have declared no competing interest.

